# Using simulated temperature regimes to test growth and development of an invasive forest insect under climate change

**DOI:** 10.1101/2021.12.06.471475

**Authors:** Jonathan A. Walter, Lily M. Thompson, Sean D. Powers, Dylan Parry, Salvatore J. Agosta, Kristine L. Grayson

## Abstract

Temperature and its impact on fitness are fundamental for understanding range shifts and population dynamics under climate change. Geographic climate heterogeneity, behavioural and physiological plasticity, and thermal adaptation to local climates makes predicting the responses of species to climate change complex. Using larvae from seven geographically distinct wild populations in the eastern United States of the non-native forest pest *Lymantria dispar dispar* (L.), we conducted a simulated reciprocal transplant experiment in environmental chambers using six custom temperature regimes representing contemporary conditions near the southern and northern extremes of the US invasion front and projections under two climate change scenarios for the year 2050. Larval growth rates increased with climate warming compared to current thermal regimes and responses differed by population. A significant population-by-treatment interaction indicated that growth rates increased more when a source population experienced the warming scenarios for their region, especially for southern populations. Our study demonstrates the utility of simulating thermal regimes under climate change in environmental chambers and emphasizes how the impacts from future increases in temperature can be heterogeneous due to geographic differences in climate-related performance among populations.

## Introduction

Climate change is altering the geographic ranges and population dynamics of organisms across the globe (Parmesan & Yohe, 2003; Thomas, 2010). Such changes reflect the accumulated effects of a shifting climate on individual fitness. Similar to other ectothermic taxa, insects are thought to be especially susceptible to direct effects of climate change due to the temperature-dependence of their vital rates (Björkman et al., 2011; Boggs, 2016); however, whether the net effects are positive or negative for population growth and viability can depend on contexts including geography, species life history, and how climate change impacts species interactions (Klapwijk et al., 2013; Van Dyck et al., 2015; Walter et al., 2018).

Numerous studies have quantified the physiological performance of various insect species in response to temperature (e.g., Kingsolver and Woods 1997, Fischer et al. 2011). However, two major caveats pertain to much of this work (Lindroth & Raffa, 2017). First, these studies often use constant-temperature thermal regimes to measure thermal performance (e.g., Thompson et al. 2017, von Schmalensee et al. 2021). Second, studies of insect response to climate warming often increase temperature in the lab or field by a constant (e.g., +2°C; Bauerfeind and Fischer 2014, Rich, et al. 2015). Given that the effects of climate change on temperature differ geographically (Karmalkar & Bradley, 2017), seasonally (Kirk et al., 2019), and diurnally (Braganza et al., 2004), such studies may provide an incomplete view of the response of insects to climate change.

These shortcomings can be overcome through the combination of modern environmental chambers and spatially downscaled climate projections including greenhouse gas forcing scenarios adopted by the Intergovernmental Panel on Climate Change (IPCC), which have relatively recently become publicly available (Eyring et al., 2016; Taylor et al., 2012). When coupled with environmental chambers capable of fine-scale temperature programming, these projections can be used to experimentally simulate temperature treatments that more accurately reflect temperature regimes found in nature. Other environmental factors potentially contributing to changes in development and fitness (e.g., light, nutrient resources, water availability, air flow) can be held constant in this type of growth chamber simulation, which allows for the effect of future temperature regimes on development and fitness to be evaluated independently. Despite its potential, however, this approach has been little used.

*Lymantria dispar dispar* (L.) (Lepidoptera: Erebidae; historical common name ‘European gypsy moth’) is an invasive forest-defoliating generalist pest in North America that feeds on over 300 host trees and causes an average of $250M USD of economic damage in the US annually (Aukema et al., 2011). Outbreaks of *L. dispar* have been implicated as a contributing factor to the decline of oaks (*Quercus* spp.) in eastern North America (Morin & Liebhold, 2016), and defoliation events alter ecosystem processes (Clark et al., 2010; Riscassi & Scanlon, 2009). Its northern invasive range limit is bounded by lethal cold temperatures for overwintering eggs and insufficient warmth to complete larval development within the shortened growing season (Gray, 2004; Streifel et al., 2019), while southern range limits may be governed by supraoptimal temperatures and a lack of sufficient chilling necessary to terminate diapause in overwintering eggs (Gray, 2004; Tobin et al., 2014). Consequently, the thermal performance of *L. dispar* under climate change may influence the location, severity, and frequency of future population outbreaks, and in turn, their cascading ecological and economic impacts.

Adding complexity to understanding the effects of climate change on *L. dispar*, recent studies have found that ecologically important traits have adapted to local climates across the invasion font. For example, egg masses sourced from warmer climates had higher viability when reared at warm range-edge temperatures relative to populations from cooler regions (Faske et al., 2019). In constant temperature experiments, warmer climate populations had lower mortality rates and smaller reductions in fitness-associated traits when reared at supraoptimal temperatures than those sourced from cooler climates (Thompson et al., 2017, 2021). Additionally, genomic evidence is consistent with a genetic basis for phenotypic differences in temperature-related performance traits (Friedline et al., 2019). The detailed knowledge on the spread and thermal performance of this invasive species makes it an ideal organism to investigate thermal performance and fitness under realistic future temperature scenarios.

This study addresses the following questions: 1) how does *L. dispar* growth and development respond to projected 2050 climatic conditions; and 2) does this response vary among populations from different parts of its invasive range? We simulated climates near the southern and northern extremes of the US invasion encompassing the larval growing season from egg hatch to pupation and measured development in seven populations. We hypothesized that climate warming would enhance fitness-related traits relative to a contemporary climate baseline under northern range-edge thermal conditions, but would reduce performance under southern range-edge conditions. Moreover, we hypothesized that the magnitude of these effects will depend on source population due to a history of local adaptation to climate, with individuals from cooler climates tending to perform better in cooler temperature regimes and individuals from warmer climates being more tolerant of warming.

## Methods

### Experimental Design

Individuals used in this experiment were sourced from seven populations in the Eastern US, a subset of those used in Thompson et al. (2021). These populations represent areas of active range expansion and the current climatic extremes of the *L. dispar* invasive range in the US (Figure 1a). Individuals used in this experiment were transported and housed under USDA APHIS permits P526P-17-03681 (KLG) and P526P-16-04388 (DP).

**Figure 1.**
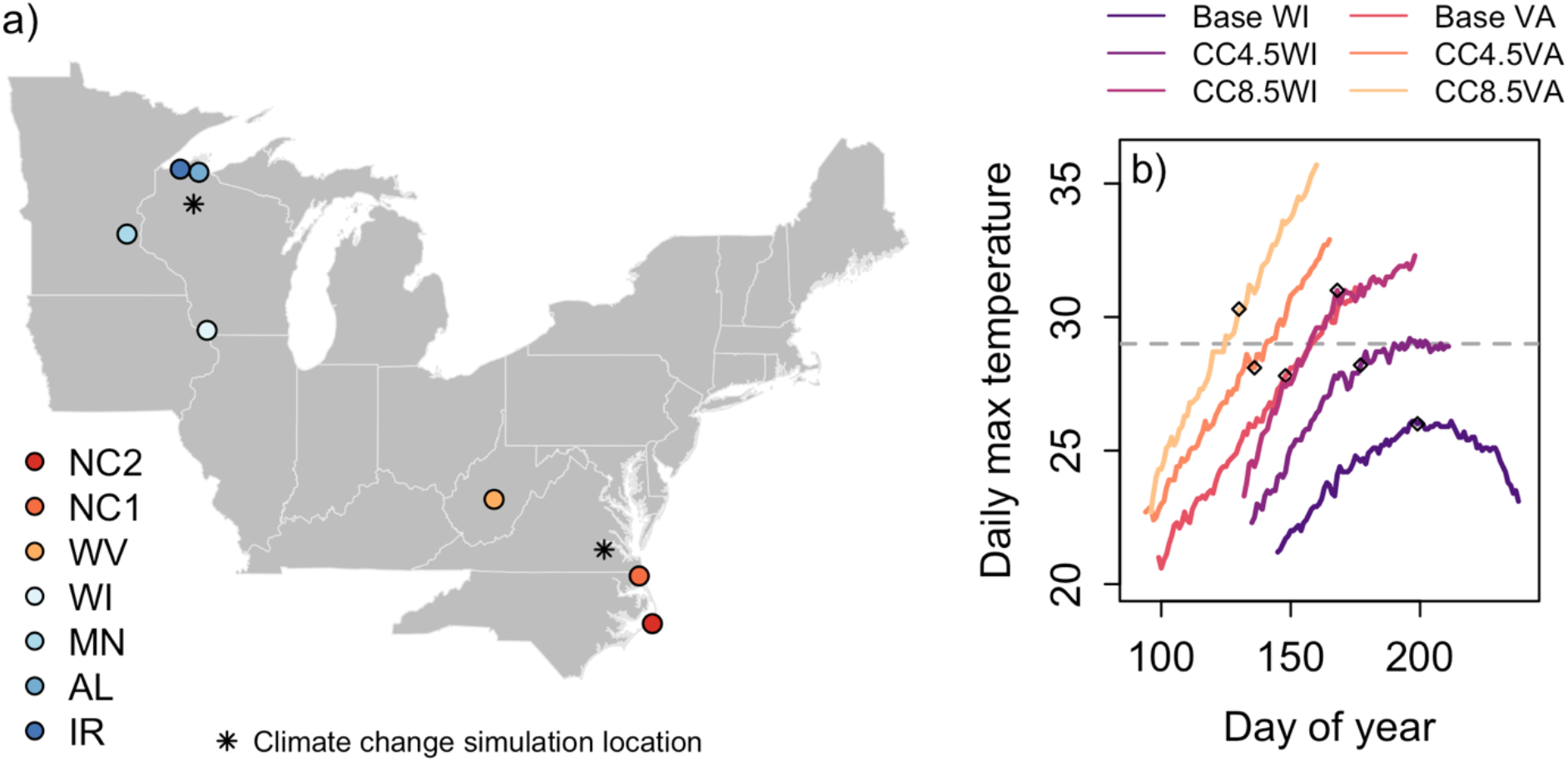
a) Map of *L. dispar* source populations. The order from south to north also corresponds to the order of mean annual temperature at each source population location; b) Daily maximum temperature over time for simulated temperature treatments, for the time period spanning modelled egg hatch through adult emergence. The grey horizontal line indicates the thermal optimum for *L. dispar* larval development (29°C). Diamonds indicate empirical 95^th^ percentile 5^th^ instar maturation dates from this experiment. Projected climate change (respectively, RCP4.5 and RCP 8.5) increased mean daily maximum temperatures during development relative to their historical baselines by 1.78°C and 4.38°C in Virginia and by 2.51°C and 4.87°C in Wisconsin. Mean daily minimum temperatures increased by 2.07°C and 4.54°C in Virginia and by 2.58°C and 4.70°C in Wisconsin.

Twenty-five individuals from each source population were reared from egg hatch to adulthood (or mortality) in six simulated thermal regimes: Baseline WI, CC4.5WI, CC8.5WI, Baseline VA, CC4.5VA, and CC8.5VA (Figure 1b). The WI and VA designations correspond to locations in Wisconsin (WI; 45.7992°N, 90.9947°W) and Virginia (VA; 37.1122°N, 77.2017°W) for which we generated thermal regimes (Figure 1a). These locations are near the inland north-western and south-eastern extremes of the invasion front and represent extremes of cold and warmth experienced by *L. dispar* in the USA. The CC4.5 and CC8.5 designations correspond to moderate and more severe representative concentration pathways, scenarios for future greenhouse gas emissions that are used to drive climate projections.

Temperature profiles were generated using BioSIM 10 software (Regniére and Saint-Amant 2017) and represent the mean of 200 replicate simulations of daily minimum and maximum temperatures. Baseline temperature regimes were based on 1981-2010 climate normals, and climate change scenarios were based on the CanRCM4 climate model (Scinocca et al., 2016). Simulated future temperature time series for each of the four nearest weather stations were constructed by adjusting 1981-2010 normals with temperature anomalies from CanRCM4 projections. We aligned the start of each chamber simulation program to the predicted date of median egg hatch as estimated using the *L. dispar* phenology model in BioSIM (Gray, 2004; Gray et al., 2001; Regniere & Sharov, 1997). The experiment began by introducing newly hatched larvae at the predicted hatch date for each simulation.

Temperature treatments were implemented in environmental chambers (Percival Scientific, Inc. model I-22VL running Intellus Connect Ultra software) on a ramp between a daily minimum temperature at 7am and a daily maximum temperature at 9pm. The chambers maintained a 14 hour light, 10 hour dark cycle with lights on between 7am and 9pm. Humidity ranged from 60-80%. The positions of individuals within the chamber were rotated to prevent micro-scale differences in air flow and light from having persistent effects on development.

Three treatments each were housed in labs at Virginia Commonwealth University (Baseline VA, Baseline WI, CC4.5WI) and at University of Richmond (CC4.5VA, CC8.5VA, CC8.5WI). Both labs used the same model of growth chamber with all settings in common. Prior to the experiment, each environmental chamber was carefully calibrated for both light and dark cycles using a two different ca. three-day programs to ensure each chamber maintained temperature within a tolerance of ±0.5°C. In the first, temperature stepped from 6°C to 16°C to 26°C. In the second, chambers were brough to a constant temperature over 8 hours before implementing two daily cycles between low and high temperatures corresponding to the beginning of the VA base temperature treatment. Temperature data loggers (HOBO U23 Pro v2, Onset Computer Corporation) were also placed inside each environmental chamber to track rearing temperatures across the experiment. These records showed that actual chamber temperatures tracked programmed temperatures across treatments, with only minor deviations that did not obscure differences among treatments (Figure S1). In addition, an earlier experiment conducted in these environmental chambers replicated two different fluctuating temperature thermal regimes in two chambers each, and found no differences in development times, masses, or survivorship between chambers implementing the same treatments (K. Grayson, *unpublished data*).

Larvae were housed in individual plastic cups with cubes of artificial diet (USDA APHIS formulation) that were replaced weekly. Individuals were checked daily between 10am and 2pm for changes in developmental stage. We recorded the following: third instar date, third instar mass, fifth instar date, fifth instar mass, pupation date, pupal mass, adult emergence date, and sex. The sex of individuals could not be determined if they died prior to the sexual dimorphism apparent in late stage larvae. We focus here on data from third and fifth instars because fungal contamination of the artificial diet increased mortality between fifth instar and adulthood in individuals raised in one of the laboratories (Figure S2). Data from this and a previous experiment (Thompson et al., 2021) showed that larval masses were strongly correlated with pupal masses (Table S1), which in turn, are an excellent proxy for fecundity (Faske et al., 2019; Honěk, 1993).

### Analyses

We analysed how growth rate from hatch to third and to fifth larval instars depended on climate treatment (Baseline WI, Baseline VA, CC4.5 WI, CC4.5 VA, CC8.5 WI, CC8.5 VA), source population (AL, IR, MN, NC1, NC2, WI1, WV1; see Figure 1), and the interaction between treatment and source population. Growth rates were computed as the difference from neonate mass divided by the development time, expressed in g day^−1^. Because individual neonate masses were below the precision of standard analytical balances, we weighed 5 replicate groups of 5 neonate larvae from each source population and took the average mass of an individual neonate larva. We included a random effect of individual sex (i.e., male, female, unknown) on the intercept because *L. dispar* become sexually dimorphic later in development. Individuals that did not survive long enough to visually determine sex (sex = unknown) were grouped together. Given that the experiments were conducted within high performance environmental chambers housed within modern climate-controlled laboratories in close geographical proximity, it is unlikely that a meaningful laboratory effect occurred, but we cannot rule it out. Analyses were conducted using linear mixed effects models with the ‘lmerTest’ package (Kuznetsova et al., 2017) in R version 3.6.1 (R Core Team, 2020). Significance of model terms was assessed using Wald *X^2^* tests with type-III sums of squares. Post-hoc comparisons between groups were considered to be significantly different based on 95% confidence intervals of estimated marginal means.

## Results

The effects of temperature regime treatment, population and treatment-by-population interaction on larval growth rates were consistent between third and fifth instars. We focus on results for fifth instars; parallel results for third instars are shown in Figure S3. Growth rates of fifth instar larvae differed by treatment (*df* = 5, *X*^2^ = 883.5, *p* < 0.0001) and population (*df* = 6, *X*^2^ = 301.0, *p* < 0.0001), with a statistically significant two-way interaction between treatment and population (*df* = 30, *X*^2^ = 89.2, *p* < 0.0001). Growth rates to fifth instar tended to increase in simulated climate change treatments (Figure 2a), and were highest in the CC4.5WI, CC8.5WI, and CC8.5VA treatments. Larvae from the WI and MN source populations tended to grow more slowly than those from other source populations (Figure 2b). The treatment-by-population interaction effect suggests that *L. dispar* larvae tend to respond to climate warming by increasing growth rate more so in simulated future climates that represented the region from where they were sourced (Figure 2c). For example, the fastest-growing populations in the CC8.5WI treatment were the two northernmost source populations, while the fastest-growing populations in the CC8.5VA treatment were the three populations from the southern range boundary.

**Figure 2.**
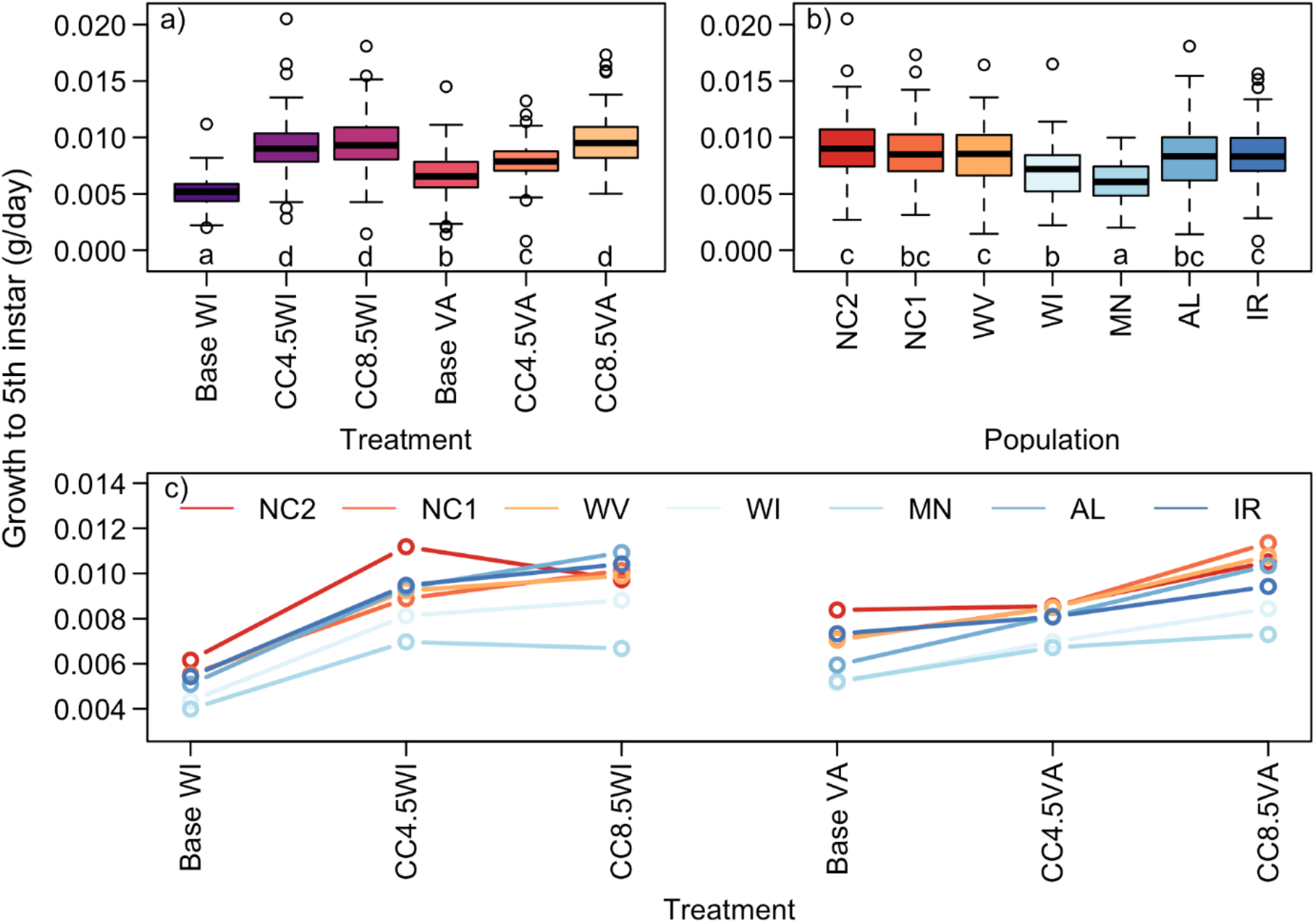
Fifth instar *L. dispar* larval masses by a) treatment, b) population, and c) population×treatment interaction for all individuals. Treatments are ordered from coolest to warmest. Populations are ordered from southernmost to northernmost. Lowercase letters in panels a) and b) denote groups whose elements have estimated marginal means with overlapping 95% confidence intervals. Sex was included in statistical models as a random effect.

## Discussion

Our study of *L. dispar* development under contemporary and future thermal regimes predicts that future larval growth rates of this destructive insect pest depend on both the degree of climate warming and geography. In general, our results suggest that larval growth rates could increase across the invasive range under climate warming based on the performance of caterpillars from different populations under simulated future thermal regimes. As hypothesized, warming increased growth rates at the northern range edge, but contrary to expectations warming did not make conditions at the southern range boundary too hot for development. Taking larval growth rates as an index of fitness due to the correlation between mass and fecundity (Faske et al., 2019) and the reduction in exposure time to natural enemies from more rapid growth, our findings suggest that, given sufficient host plant resources, future changes in fitness will depend on the population location and magnitude of temperature change. If climate warming causes geographically dependent trends in *L. dispar* fitness, there would be substantial effects on the future ecological and economic impacts of this forest pest, and on allocation of management efforts to slow its spread.

We found evidence that source populations responded differently to experimental treatments (Figure 2c). However, evidence for local adaptation of larval growth and development was equivocal overall, in contrast to other recent studies (Faske et al., 2019; Friedline et al., 2019; Thompson et al., 2017, 2021). Among-population variation in larval growth rates in the baseline and CC4.5 treatments was not discernibly related to a “home field advantage” as was seen in the CC8.5 treatments. One possibility is that only in the CC8.5 treatments did supraoptimal temperatures occur frequently enough during larval development for growth rates to diverge. Studies testing thermal performance under constant temperatures may be more likely to see population variation in responses to thermal extremes, while this experiment used fluctuating regimes in which supraoptimal temperatures were transient, even under climate change treatments. There is evidence that southern populations are more tolerant of high temperatures (Thompson et al., 2017, 2021), which could explain their better performance in our hottest treatments. However, we found no clear evidence that northern populations can perform better than southern populations in a colder climate (Figure 2c). Overall, these results continue to demonstrate that while larval growth and development in *L. dispar* has undergone climate-related adaptation, these traits remain plastic in response to temperature (Thompson et al. 2021).

Future *L. dispar* performance with climate warming could make range expansion and population outbreaks more common in the northern range extremes. Indeed, establishment of populations in northern Minnesota has occurred in areas predicted to be marginal for survival (Streifel et al., 2019). Conversely, at the southern extreme, range stasis and retraction has already been observed (Tobin et al., 2014) but our results suggest that larval development at the southern range edge is not impaired by present or projected future temperature regimes. Reduced egg viability, possibly due to insufficient cold to complete diapause, could be the mechanism for range stasis at the southern extreme (Faske et al., 2019; Gray, 2004). More broadly, our predicted effects could be amplified or negated by effects of climate change on life stages not considered in this study (Kingsolver & Buckley, 2020), or by effects on ecological relationships, e.g., with host plants or natural enemies. Geographical changes in the propensity for range expansion and population outbreaks would be of considerable concern to extensive management efforts to slow spread and protect land from damaging outbreaks (Tobin et al., 2012).

The ability to test experimental thermal regimes featuring spatially explicit climate change projections was an innovative approach that greatly enhanced the realism of this study while retaining the ability afforded by environmental chambers to maintain consistency in unmanipulated conditions. As climate change has non-uniform effects on the spatiotemporal distributions of temperatures (Braganza et al., 2004; Karmalkar & Bradley, 2017; Kirk et al., 2019), our approach represents a substantial advance in methodology compared to simply increasing temperature by a constant value from a current baseline. Additionally, programmed environmental chambers provide the ability to test realistic thermal regimes from any geographic position or point in time independent of physical location. We are unaware of other studies taking this approach, likely in part because its feasibility depends on modern environmental chamber control software and a study system with some *a priori* knowledge of phenology, but we encourage this design as a means of increasing the realism of climate change ecophysiology studies in controlled experimental settings.

A robust body of literature on insects has shown that warmer temperatures can facilitate range expansion (Lehmann et al., 2020) or can accelerate invasion speed (Seiter & Kingsolver, 2013), but may also negatively impact populations of other species (Haynes et al., 2014; Johnson et al., 2010; Klapwijk et al., 2013). Such variations in thermal responses impede generalization of the response of species to climate change. Our findings, taken together with other studies documenting geographical variation in thermal response of *L. dispar* (Faske et al., 2019; Thompson et al., 2017, 2021), emphasize how the ecological effects of climate change can be spatially heterogeneous, not only due to regional variation in climate change (Karmalkar & Bradley, 2017), but also due to variation in the thermal tolerance and performance of local populations of a given species.

## Acknowledgements

This work was supported by the National Science Foundation (DEB 1702701), the Slow the Spread Foundation (DP and KG), the University of Richmond School of Arts & Sciences, and the Virginia Commonwealth University Center for Environmental Studies. Remi Saint-Amant generated temperature time series. T. Faske, C. Jahant-Miller, K. Onufrieva, C. Foelker, K. Theilen-Cremers, C. Elder, J. Johnson, S. Hoffman, C.J. Campbell and V. Huelsman helped acquire source material for this study. S. Goetz and C. Jahant-Miller assisted in maintaining population cultures. K. Sanchez, R. Ostrom, C. Miller, P. Hafker, A. Burnett, L. Milner, and P. Gibbs helped rear larvae and collect data.

## Online Supplement

**Table S1.** Correlations between larval and pupal masses. Correlations are separated by sex due to sexual dimorphism. Data are taken from this study and from Thompson et al. (2021), which used constant temperature thermal regimes.

| **Study** | **Instar** | **Sex** | ***n*** | **$\hat{\rho}$** | ***p*** |
| --- | --- | --- | --- | --- | --- |
| This study | 3<sup>rd</sup> | M | 358 | 0.11 | 0.037 |
| This study | 3<sup>rd</sup> | F | 272 | 0.07 | 0.075 |
| This study | 5<sup>th</sup> | M | 354 | 0.30 | <0.001 |
| This study | 5<sup>th</sup> | F | 266 | 0.37 | <0.001 |
| Thompson et al. (2021) | 3<sup>rd</sup> | M | 557 | 0.37 | <0.001 |
| Thompson et al. (2021) | 3<sup>rd</sup> | F | 475 | 0.13 | 0.004 |

**Figure S1.**
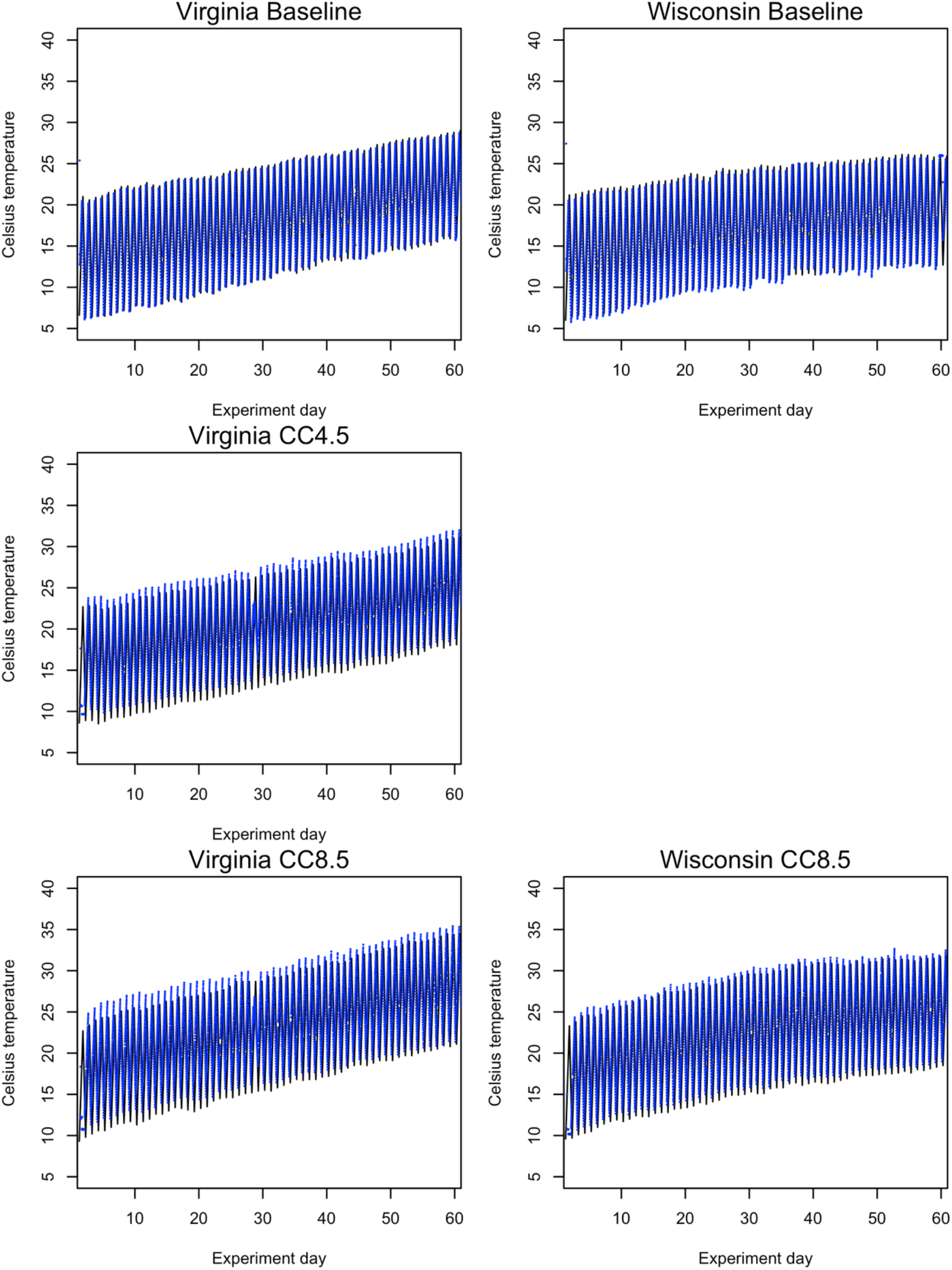
Comparison between environmental chamber programmed temperature (black lines) and recorded temperature (blue points). Wisconsin CC4.5 is not shown because the sensor failed to record data. By day 60, every surviving larva had reached 5^th^ instar so the x-axis is limited to days 1 through 60.

**Figure S2.**
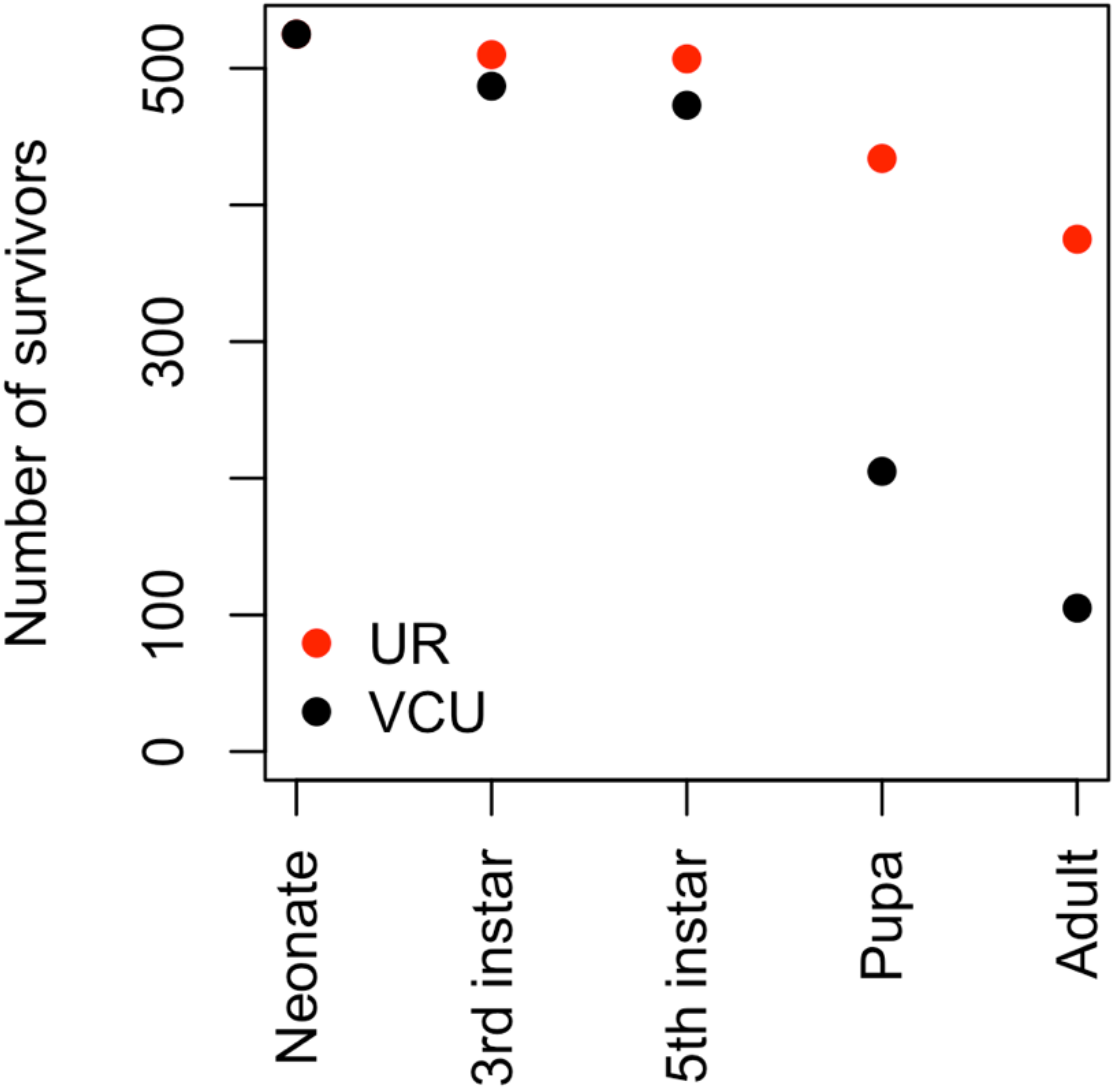
Number of survivors by life stage and laboratory. Larvae were introduced into environmental chambers as neonates and monitored for development into 3^rd^ and 5^th^ larval instars, pupae, and adults. A mold outbreak in the VCU laboratory caused apparent reductions in survival between 5^th^ instar and pupation, analyses presented in this manuscript focus on data for larvae.

**Figure S3.**
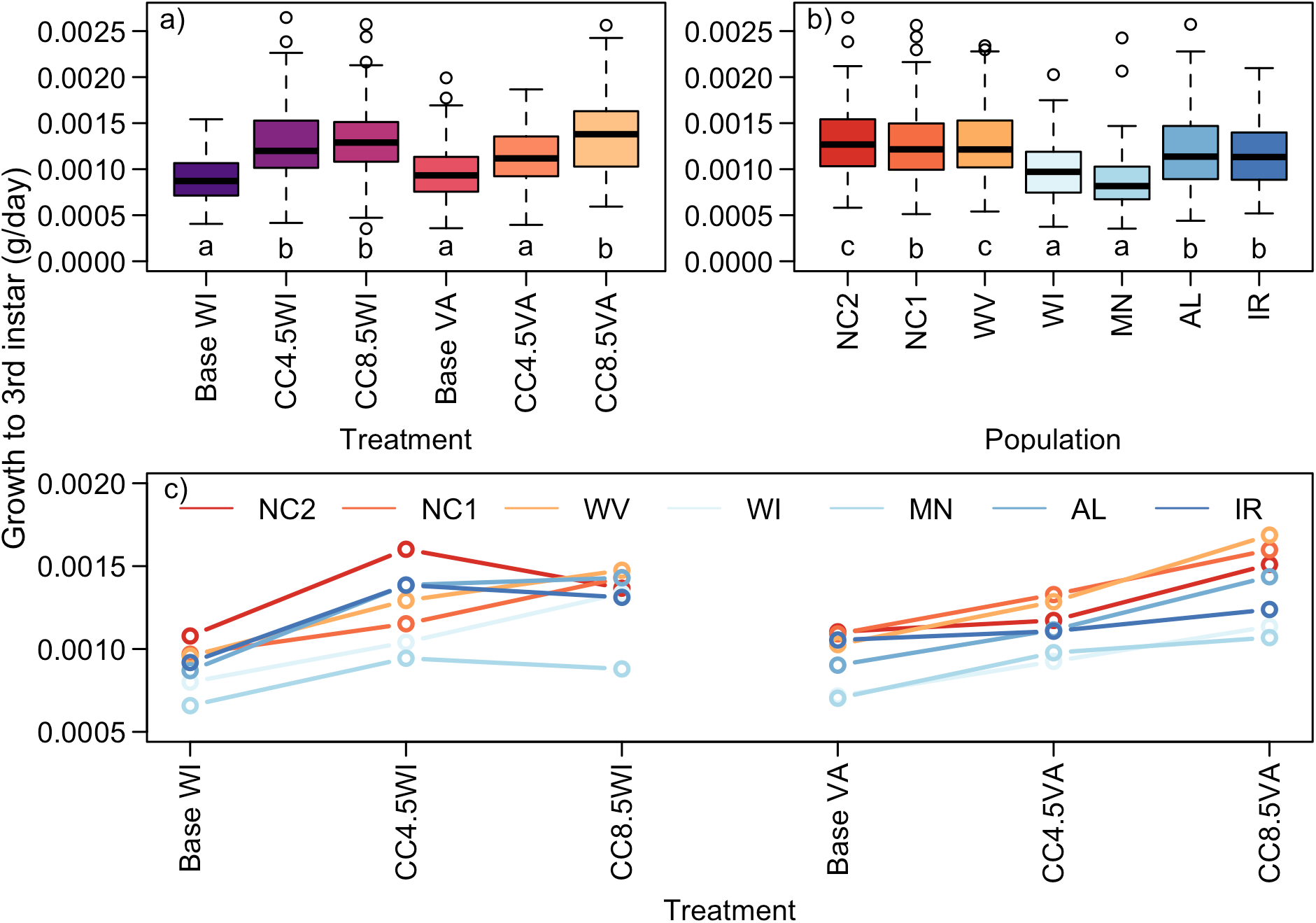
Third instar larval masses by a) treatment (df = 5, *X*^2^ = 349.47, *p* < 0.0001), b) population (df = 6, *X*^2^ = 251.031, *p* < 0.0001), and c) population×treatment interaction (df = 30, *X*^2^ = 92.16, *p* < 0.0001)for all individuals. Treatments are ordered from coolest to warmest. Populations are ordered from southernmost to northernmost. Lowercase letters in panels a) and b) denote groups whose elements have estimated marginal means with overlapping 95% confidence intervals. Sex was included in statistical models as a random effect.

